# Benchmarking single-cell foundation models reveals an advantage in the low-data regime for cell-type and disease-state prediction

**DOI:** 10.64898/2026.10.12.758421

**Authors:** Haya Benhayon, Michael M. Danziger, Danny Ben-Zvi, Michal Rosen-Zvi

## Abstract

Single-cell foundation models perform well on cell-type classification, but their value for disease-state prediction and their advantage over classical representations remains unclear. We benchmarked cell-representations from four foundation models using logistic regression, on 868 disease-prediction tasks (82 datasets) and 65 cell-type datasets from CELLxGENE. With full data, the baseline (lognormalized counts) outperformed the best model (TranscriptFormer-Sapiens) for cell type and matched it for disease. However, with few labeled cells, TranscriptFormer-Sapiens significantly outperformed the baseline in both tasks up to ∼ **10^2^** samples per class. Our comprehensive zero-shot benchmark identifies the data-scarce regime as a domain where Single-cell foundation models truly show advantage.

## 1 Background

Large-scale pretraining is a powerful strategy for learning transferable representations across domains, most notably in natural language processing models [1]. Their strength lies in capturing complex, high-order patterns that are difficult to detect using previous approaches. Similar architectures have been adapted to single-cell RNA-seq (scRNA-seq) analyses, where models are trained on millions of cells to learn transferable representations of gene expression. Single-cell foundation models (scFMs) have demonstrated strong performance in established tasks such as cell-type classification [2, 3], but their value for more complex downstream tasks remains unclear. Diseasestate detection is of central importance for biomedical research and clinical utility. However, whereas cell identity is readily inferred from transcriptomic profiles, disease-associated transcriptional signals are more heterogeneous and context dependent, and results have been mixed. Rigorous and comprehensive evaluation of biomedical foundation models remains a challenge [4], and evaluation of scFMs for disease-state classification has been limited [5–7].

Several recent works have raised questions about the zero-shot performance of scFMs. Boiarsky et al. [8] found that L1-regularized logistic regression matched or outperformed scBERT for cell-type annotation in two datasets. Kedzierska et al. [9] found frozen Geneformer and scGPT embeddings were outperformed by highly variable gene selection, Harmony and scVI in cell-type clustering and batch integration across five datasets, and Souza and Mehta [10] found that scTOP, a parameter-free linear projection, matches or exceeds scFMs in cross-species annotation, human cell-type classification, and in a single-disease setting.

## 2 Results and Discussion

We set out to systematically evaluate the advantages that scFMs for scRNA-seq, namely scVI [11], Geneformer [3], TranscriptFormer-Sapiens (TF-SAPIENS, the human-only checkpoint) [6], and BMFM-RNA [12] may provide over classical approaches in disease-associated prediction tasks, and if those advantages are impacted by the number of samples available for learning. Using human scRNA-seq data from CellxGene Census (version 2025-01-30) [13], we benchmarked classical representations: log-normalized counts and PCA-reduced features, against embeddings from these four foundation models using a logistic regression classifier, yielding 868 disease-prediction tasks across 82 datasets and 65 datasets for the cell-type benchmark (see Methods) (Fig. 1). We used donor-stratified cross-validation, keeping all cells from each donor in either the training or test fold, to prevent classifiers from exploiting donor-specific effects. Disease-prediction was evaluated within each cell-type independently, and the benchmark corpus was restricted to studies with multiple diseases for the cell type task also, to avoid selection bias when comparing disease and cell type performance. Methods were compared in three regimes: full-data training, low-data conditions, and median-pooling aggregation in which each disease-donor-celltype-dataset sample is represented as the median of the representation vectors for all cells sharing these labels. For raw samples, median-pooling aggregation is equivalent to median pseudobulk, the recommended method for log-normalized data.

**Fig 1:**
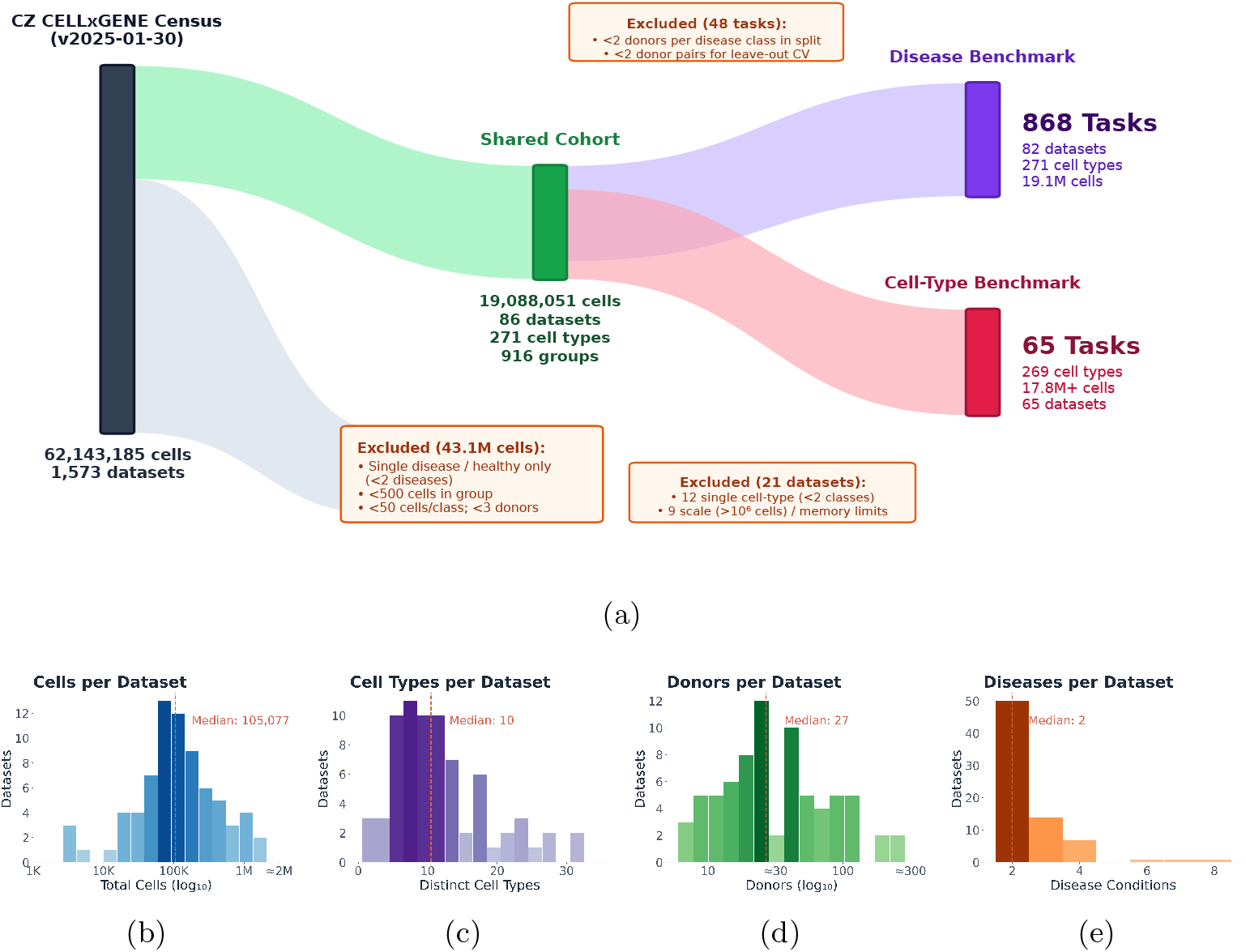
Data curation pipeline and cohort overview. **(a)** Quantitative inclusion and exclusion flow diagram from CZ CELLxGENE Census down to the disease-prediction and cell-type prediction benchmark cohorts. **(b)** Distribution of total cells per dataset (log_10_ scale). **(c)** Distribution of distinct cell types per dataset. **(d)** Distribution of donors per dataset (log_10_ scale). **(e)** Distribution of disease conditions per dataset.

In cell-type prediction, Raw log-normalized counts achieved the highest absolute F1-macro score, with TF-SAPIENS and BMFM-CONCAT close behind (Fig. 2c Supplementary Table S1). Despite this small absolute gap, Raw log-norm ranked significantly ahead of TF-SAPIENS and all other representations by Friedman analysis [14] (Fig. 2a and Supplementary Fig. S1a). On Matthews correlation coefficient (MCC) [15], Raw log-norm and TF-SAPIENS achieved identical absolute scores and were statistically indistinguishable, although both ranked significantly ahead of the remaining representations (Fig. 2a Supplementary Table S1). In the low-data regime, scFMs showed a significant advantage over classical representations: TF-SAPIENS outperformed Raw Log-norm on 100% of cell-type datasets at *n* = 2 and *n* = 4 cells per class (Fig. 2b and Supplementary Fig. S2b). Beyond 64 cells per class, scFM did not confer an advantage.

**Fig 2:**
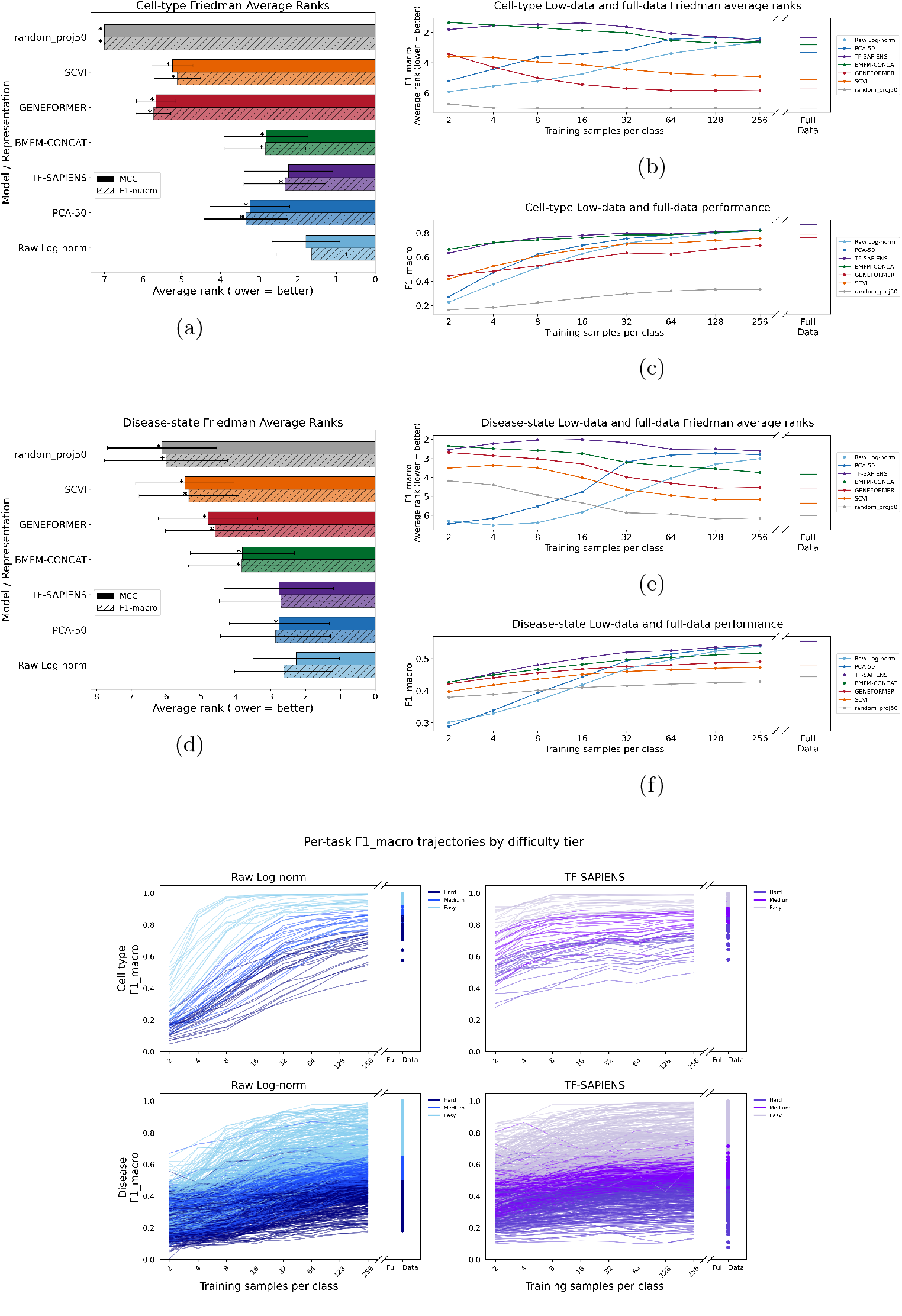
Cell-type and disease-state prediction benchmarks. (a) Friedman average ranks (lower = better) across all representations under full-data training prediction, 5(MCC and F1-macro), cell-type *T* = 65 datasets. asterisks mark representations significantly different from Raw Log-norm (pairwise Wilcoxon signed-rank, Holm-corrected, *p <* 0.05; full pairwise comparisons in Supplementary Fig S1a). (b) Friedman average ranks (F1-macro only) across the low-data, cell-type prediction, with full-data ranks shown at right for comparison. (c) Mean F1-macro per representation, averaged across dataset-level means, across the low-data, cell-type prediction and at full data. (d) Friedman average ranks under full-data training (MCC and F1-macro), disease prediction, dataset-level correction applied (*T* = 82 datasets); asterisks as in (a). (e) Friedman average ranks (F1-macro only) across the low-data, disease prediction, using dataset-collapsed significance testing throughout (see Methods), with full-data ranks shown at right. (f) Mean F1-macro per representation, averaged across dataset-level means, across the low-data, disease prediction and at full data. (g) Per-task F1-macro trajectories across the low-data sweep for Raw Log-norm and TF-SAPIENS, colored by difficulty tier (Hard/Medium/Easy), cell-type (top row) and disease (bottom row); full-data values shown at right of each panel. Error bars in (b,c,e,f) omitted for clarity.

Disease prediction proved substantially more challenging, with greater variability across datasets and methods (Fig. 2f Supplementary Table S2). Under full-data training, Raw log-norm, TF-SAPIENS, and PCA-50 achieved nearly identical absolute performance on both metrics and, as a group, ranked significantly ahead of all other foundation models. Within this top group, the significance pattern differed by metric: on MCC, Raw log-norm ranked significantly ahead of PCA-50, yet neither differed significantly from TF-SAPIENS; on F1-macro, all three were statistically indistinguishable (Supplementary Fig. S1b). In the low-data regime, TF-SAPIENS consistently outperformed Raw Log-norm at small sample sizes. The TF-SAPIENS advantage over Raw Log-norm on F1-macro remained significant through 128 cells per class (Fig. 2e).

To determine whether the scFM advantage depends on task difficulty, we stratified tasks into three equal-sized tiers (Hard, Medium, and Easy) based on full-data classical baselines performance (see Methods), and compared TF-SAPIENS to Raw log-norm. Similar trends were observed across the three difficulty tiers, with TF-SAPIENS out-performing Raw log-norm in low data regimes, with no difference between the models when data is abundant.(Supplementary Fig. S2a). Together, these results suggest that the principal advantage of current scFMs at zero-shot is improved sample efficiency rather than superior performance when training data are abundant.

Median-pooling aggregation showed similar results. However, BMFM-RNA emerged as significantly best for cell-type prediction (dataset-level Friedman vs. TF-SAPIENS), whereas for disease prediction there was no difference between scFMs and classical representations. Median pooling, which discards per-cell heterogeneity, sharpens representational differences for cell-type prediction but erodes them for disease prediction (Supplementary Fig. S1c–f).

Our study is limited by the quality of the CellXGene corpus, where cell-type or disease labels may not be of uniform quality. We used a logistic regression classification for its simplicity and relative lack of hyperparameters. We further limited our perspective to the representation quality itself (zero-shot), as it may be the simplest use case. Recent models including experimentally grounded supervision, as in AlphaGenome [16], may provide higher quality representations. The robustness of our findings under fine-tuning or with different predictive models is an important topic for future research.

## 3 Conclusion

Claims that scFMs outperform classical methods have been challenged by reports that simple baselines match them. Our systematic evaluation, across 868 disease-prediction tasks and 65 cell-type datasets, reconciles these views: zero-shot foundation-model embeddings do outperform classical representations, but only when labeled data are scarce, with a significant advantage up to about *∼* 10^2^ cells per class. With abundant labels, log-normalized counts matched or outperformed every model. We therefore find that foundation models are most valuable for rare cell populations and small cohorts.

## 4. Methods

Candidate tasks were derived systematically from the CZ CELLxGENE Census (version 2025-01-30) [13]. To eliminate selection bias when comparing disease-state and cell-type classification, both benchmarks were evaluated on as close as possible to a single shared corpus, applying objective sample-size criteria across the entire Census rather than selectively sampling favorable datasets (Fig 1a). Because disease prediction requires contrasting clinical states, single-condition and healthy-only studies were excluded, which accounts for the vast majority of excluded Census cells. The remaining corpus was filtered to (dataset, cell type) groups with *≥*500 cells, *≥* 2 disease conditions (*≥* 50 cells each), and *≥* 3 donors per disease stratum to support donor-stratified cross-validation.

From this shared foundation: for disease prediction, tasks were evaluated per cell type; 48 of 916 candidate groups were excluded because their donor distributions could not support valid cross-validation splits (leaving *<* 2 donors for a disease class in a fold), yielding 868 evaluated tasks across 82 datasets. For cell-type prediction, tasks were evaluated per dataset, requiring *≥* 2 qualifying cell types (12 single-cell-type datasets excluded) and omitting 9 datasets (*>* 10^6^ cells) that exceeded memory limits during foundation model embedding extraction, yielding 65 evaluated datasets across 269 cell types.

All representations were evaluated with L2-regularized logistic regression (*α* = 10^−5^) as a common classifier. Classical representations were log-normalized expression (Raw log-norm), PCA using 50 components, fit on the training fold and applied to the test fold to avoid leakage (PCA-50), and a random 50-dimensional projection as a lower-bound control (random proj50). FM representations came from scVI [11], Geneformer [3] and TranscriptFormer-Sapiens [6] as exposed via the CellXGene Census API; BMFM-RNA embeddings (BMFM-CONCAT) concatenated the CLS token representations from the WCED Multitask and MLM Multitask (v2 LLaMA) check-points as described in [12] and in the released code for this project. All methods used donor-stratified *k*-fold cross-validation (*k* = 5, limited by the minimum donor count per disease stratum; with additional disease stratification for cell-type prediction).

Three regimes were evaluated: (i) full-data training; (ii) low-data subsampling to 2, 4, 8, 16, 32, 64, 128 and 256 cells per class (5 bootstrap resamples per size); and (iii) median-pooling aggregation, taking the median expression or embedding value per donor (disease) or per donor and cell type (cell-type) before classification.

Performance was measured by macro-averaged F1 [17] and the Matthews correlation coefficient (MCC) [15], both informative under the class imbalance typical of disease-prediction tasks [18]. For a confusion matrix *C* over *K* classes, let *c* = *Σ*_*k*_ *C*_*kk*_, s = *Σ*_*k,l*_ *C*_*kl*_, *t*_*k*_ = *Σ*_*l*_ *C*_kl_ and _*pk*_= Σ_*l*_ *C*_*lk*_ (correct, total, true and predicted counts). MCC was computed as

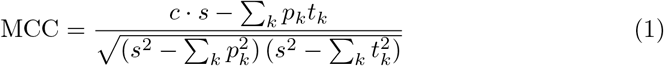

and macro-averaged F1 as the unweighted mean of per-class F1 scores:

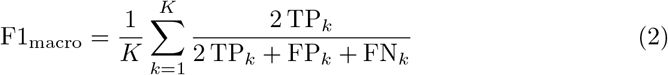

where TP_*k*_, FP_*k*_ and FN_*k*_ are the per-class true positives, false positives and false negatives, *K* is the number of classes present in the true or predicted test-fold labels, and per-class F1 is 0 when undefined. Metrics were computed per fold on cell-level predictions (excluding test cells whose label was absent from the training fold) and averaged across folds.

Representations were compared by the Friedman test (Iman-Davenport correction) on average ranks, with pairwise Wilcoxon signed-rank tests and Holm correction as post-hoc comparisons [19, 20]. For disease prediction, significance was assessed on dataset-collapsed scores (mean per dataset rather than per task) to account for clustering of tasks within datasets. Cell-type tasks were evaluated separately per dataset and so needed no correction.

Difficulty tiers (Hard, Medium, Easy) were tertiles of the mean full-data F1-macro of Raw Log-norm and PCA-50. Within each tier, TF-SAPIENS and Raw Log-norm were compared by one-sided paired Wilcoxon signed-rank tests (TF-SAPIENS *>* Raw Log-norm) at each low-data size, with Holm correction across the eight sizes within each benchmark-tier combination, using dataset-collapsed scores for disease.

## Declarations

### Ethics approval and consent to participate

Not applicable.

### Consent for publication

Not applicable.

### Availability of data and materials

All single-cell RNA-sequencing data are publicly available through the CZ CELLx-GENE Census (release 2025-01-30) [13]. All code used for data retrieval, benchmarking, statistical analysis, and figure generation is available at [21].

## Acknowledgments

We thank members of our groups for fruitful discussion.

