## Supplementary Information for "Benchmarking single-cell foundation models reveals an advantage in the low-data regime for cell-type and disease-state prediction"

[1]Department of Developmental Biology and Cancer Research, The Institute for Medical Research Israel-Canada, Jerusalem, Israel.
[2]IBM Research.
[3]Center for Computational Medicine, The Hebrew University of Jerusalem, Jerusalem, Israel.
[4]Faculty of Medicine, The Hebrew University of Jerusalem, Jerusalem, Israel.
[5]Merck Group.

Contributing authors:;

### Supplementary Tables

**Table S1**: Cell-type: Full-data. Values are mean ± standard deviation.

| Metric | Raw Log-norm | PCA-50 | TF-SAPIENS | BMFM-CONCAT | GENEFORMER | SCVI | Random Proj.-50 |
|---|---|---|---|---|---|---|---|
| **MCC** | 0.898 ± 0.085 | 0.878 ± 0.112 | 0.898 ± 0.085 | 0.894 ± 0.084 | 0.807 ± 0.133 | 0.826 ± 0.140 | 0.488 ± 0.207 |
| **F1 Macro** | 0.871 ± 0.098 | 0.841 ± 0.131 | 0.867 ± 0.103 | 0.866 ± 0.098 | 0.763 ± 0.144 | 0.788 ± 0.151 | 0.443 ± 0.192 |
| **Recall Macro** | 0.873 ± 0.096 | 0.857 ± 0.118 | 0.885 ± 0.090 | 0.882 ± 0.088 | 0.792 ± 0.133 | 0.818 ± 0.137 | 0.515 ± 0.195 |
| **Balanced Accuracy** | 0.873 ± 0.096 | 0.857 ± 0.118 | 0.885 ± 0.090 | 0.882 ± 0.088 | 0.792 ± 0.133 | 0.818 ± 0.137 | 0.515 ± 0.195 |

**Table S2**: Disease-state: Full-data. Values are mean ± standard deviation.

| Metric | Raw Log-norm | PCA-50 | TF-SAPIENS | BMFM-CONCAT | GENEFORMER | SCVI | Random Proj.-50 |
|---|---|---|---|---|---|---|---|
| **MCC** | 0.345 ± 0.275 | 0.339 ± 0.279 | 0.333 ± 0.279 | 0.295 ± 0.257 | 0.219 ± 0.225 | 0.176 ± 0.206 | 0.099 ± 0.129 |
| **F1 Macro** | 0.583 ± 0.175 | 0.582 ± 0.175 | 0.581 ± 0.175 | 0.560 ± 0.161 | 0.519 ± 0.147 | 0.491 ± 0.144 | 0.448 ± 0.119 |
| **Recall Macro** | 0.629 ± 0.169 | 0.631 ± 0.170 | 0.633 ± 0.172 | 0.612 ± 0.159 | 0.572 ± 0.150 | 0.544 ± 0.149 | 0.500 ± 0.128 |
| **Balanced Accuracy** | 0.643 ± 0.160 | 0.646 ± 0.162 | 0.648 ± 0.164 | 0.627 ± 0.152 | 0.587 ± 0.143 | 0.557 ± 0.141 | 0.511 ± 0.116 |

**Table S3**: Cell-type: Median-pooling. Values are mean ± standard deviation.

| Metric | Raw Log-norm | PCA-50 | TF-SAPIENS | BMFM-CONCAT | GENEFORMER | SCVI | Random Proj.-50 |
|---|---|---|---|---|---|---|---|
| **MCC** | 0.913 ± 0.075 | 0.921 ± 0.075 | 0.921 ± 0.086 | 0.938 ± 0.068 | 0.863 ± 0.124 | 0.892 ± 0.092 | 0.816 ± 0.136 |
| **F1 Macro** | 0.913 ± 0.079 | 0.923 ± 0.075 | 0.921 ± 0.087 | 0.938 ± 0.070 | 0.860 ± 0.127 | 0.890 ± 0.096 | 0.820 ± 0.136 |
| **Recall Macro** | 0.917 ± 0.073 | 0.926 ± 0.072 | 0.925 ± 0.083 | 0.940 ± 0.065 | 0.870 ± 0.118 | 0.897 ± 0.089 | 0.827 ± 0.131 |
| **Balanced Accuracy** | 0.917 ± 0.073 | 0.926 ± 0.072 | 0.925 ± 0.083 | 0.940 ± 0.065 | 0.870 ± 0.118 | 0.897 ± 0.089 | 0.827 ± 0.131 |

**Table S4**: Disease-state: Median-pooling. Values are mean ± standard deviation.

| Metric | Raw Log-norm | PCA-50 | TF-SAPIENS | BMFM-CONCAT | GENEFORMER | SCVI | Random Proj.-50 |
|---|---|---|---|---|---|---|---|
| **MCC** | 0.333 ± 0.293 | 0.366 ± 0.305 | 0.382 ± 0.320 | 0.378 ± 0.310 | 0.338 ± 0.337 | 0.301 ± 0.324 | 0.278 ± 0.307 |
| **F1 Macro** | 0.567 ± 0.183 | 0.598 ± 0.185 | 0.603 ± 0.195 | 0.601 ± 0.186 | 0.583 ± 0.195 | 0.562 ± 0.191 | 0.549 ± 0.183 |
| **Recall Macro** | 0.617 ± 0.168 | 0.647 ± 0.177 | 0.662 ± 0.186 | 0.659 ± 0.182 | 0.639 ± 0.192 | 0.609 ± 0.189 | 0.597 ± 0.184 |
| **Balanced Accuracy** | 0.621 ± 0.165 | 0.653 ± 0.173 | 0.669 ± 0.181 | 0.668 ± 0.177 | 0.646 ± 0.189 | 0.615 ± 0.184 | 0.604 ± 0.179 |

**Table S5**: **Tasks below the target low-data sample size.** Number and percentage of cell-type and disease-state prediction tasks where at least one class had fewer cells available than the requested sample size. the task was still evaluated with the available cells.

**A. Cell-type prediction**

| Cells/class | Tasks | Below target | Below target (%) |
|---|---|---|---|
| 2 | 65 | 0 | 0.0 |
| 4 | 65 | 0 | 0.0 |
| 8 | 65 | 0 | 0.0 |
| 16 | 65 | 0 | 0.0 |
| 32 | 65 | 0 | 0.0 |
| 64 | 65 | 0 | 0.0 |
| 128 | 65 | 1 | 1.6 |
| 256 | 65 | 2 | 3.2 |

**B. Disease-state prediction**

| Cells/class | Tasks | Below target | Below target (%) |
|---|---|---|---|
| 2 | 868 | 0 | 0.0 |
| 4 | 868 | 0 | 0.0 |
| 8 | 868 | 0 | 0.0 |
| 16 | 868 | 0 | 0.0 |
| 32 | 868 | 0 | 0.0 |
| 64 | 868 | 9 | 1.0 |
| 128 | 868 | 82 | 9.5 |
| 256 | 868 | 284 | 32.9 |

#### Supplementary Figures

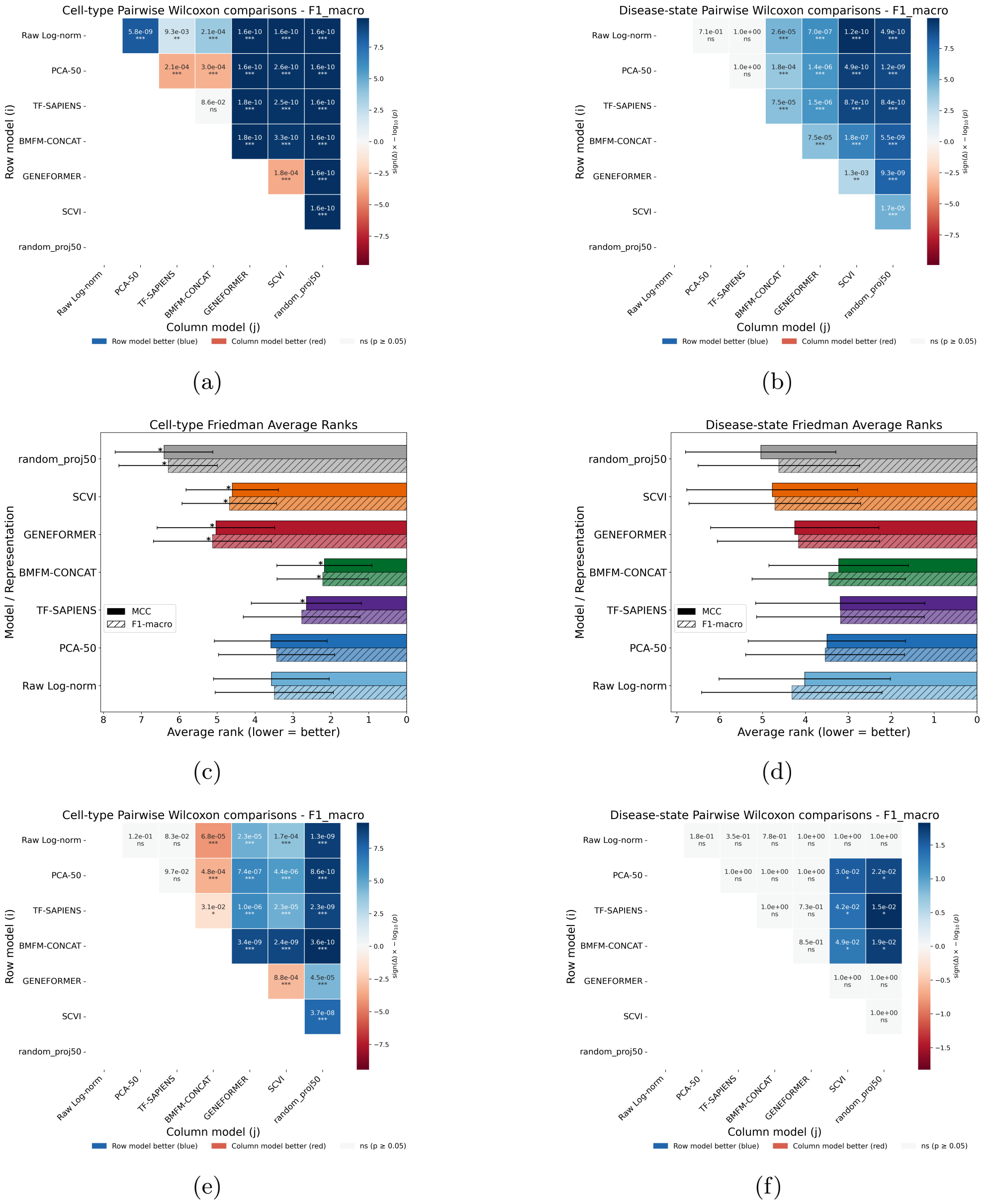


**Fig. S1**: (a,b) Pairwise comparison of representations under full-data training (F1-macro) for cell-type prediction (a: $T = 65$ datasets) and disease-state prediction (b: $T = 82$ datasets, dataset-collapsed). Each cell compares a row and a column representation using a paired Wilcoxon signed-rank test with Holm correction over all 21 pairs; color indicates which representation has the higher mean F1-macro, and marked cells indicate $p_{\mathrm{Holm}} < 0.05$. (c) Cell-type prediction under median-pooling aggregation, MCC and F1 with Friedman-ranked standard deviation across methods. (d) Disease-state prediction under median-pooling aggregation, MCC and F1 with Friedman-ranked standard deviation across methods. (e,f) Pairwise comparison of representations under median-pooling aggregation (F1-macro) for cell-type prediction (e: $T = 65$ datasets) and disease-state prediction (f: $T = 82$ datasets, dataset-collapsed). Each cell compares a row and a column representation using a paired Wilcoxon signed-rank test with Holm correction color indicates which representation has the higher mean F1-macro, and marked cells indicate $p_{\mathrm{Holm}} < 0.05$.

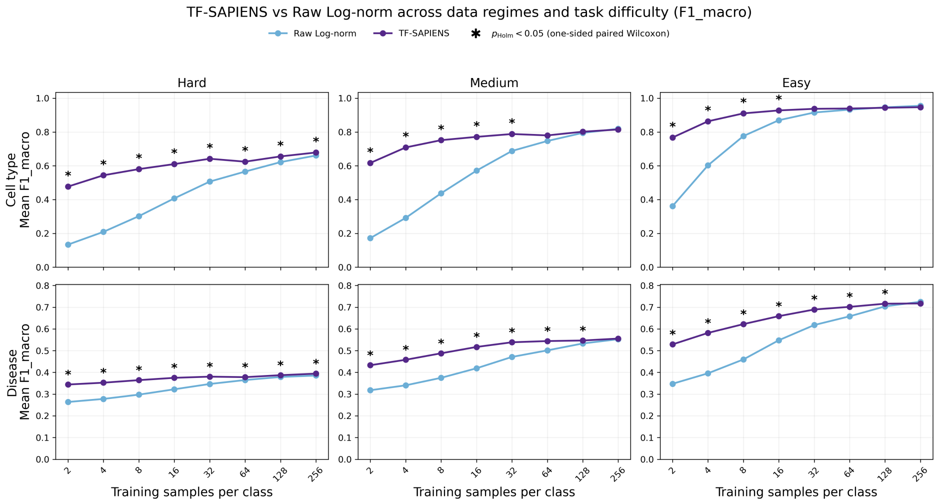


(a)

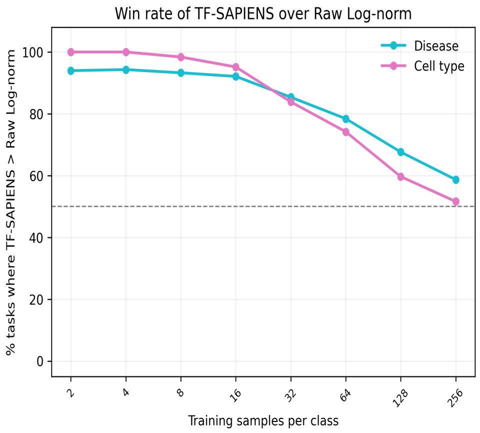


(b)

**Fig. S2**: (a) TF-SAPIENS advantage over Raw Log-norm by task difficulty tier (Hard/Medium/Easy). Mean F1-macro of TF-SAPIENS and Raw Log-norm across the low-data sweep, by benchmark (cell type, top row; disease, bottom row) and tier. Disease scores were collapsed to one value per dataset before testing. Asterisks mark sample sizes $n$ at which TF-SAPIENS is significantly higher than Raw Log-norm (one-sided paired Wilcoxon signed-rank test, Holm-corrected across sample sizes within each panel, $p < 0.05$). (b) Head-to-head comparison of raw log-normalized counts versus TF-SAPIENS, expressed as percent F1-macro.
